# Targeting REV7 exposes a mitotic vulnerability in colorectal cancer cells

**DOI:** 10.64898/2026.09.29.755408

**Authors:** Madison Guyette, Sombodhi Bhattacharya, Naveen Kothandaraman, Vihit Gupta, Andrew Crompton, Katherine Boyle, Noah Harrahill, Jon Ramsey, Dmitry M. Korzhnev, M. Kyle Hadden, Nimrat Chatterjee

**Affiliations:** Department of Microbiology and Molecular Genetics, University of Vermont, Burlington, Vermont, U.S.A. 05405; Department of Pharmaceutical Sciences, University of Connecticut, Storrs, Connecticut, U.S.A. 06269; Larner College of Medicine, University of Vermont, Burlington, Vermont, U.S.A. 05405; Department of Biochemistry, University of Vermont, Burlington, Vermont, U.S.A. 05405; Department of Molecular Biology and Biophysics, University of Connecticut Health Center, Farmington, Connecticut, U.S.A 06030; University of Vermont Cancer Center, University of Vermont, Burlington, Vermont, U.S.A. 05405

**Keywords:** REV7, translesion synthesis, colorectal cancer, therapeutic resistance, mitotic disruption, chromosomal instability, micronuclei, colchicine

## Abstract

Therapeutic resistance remains a barrier to durable treatment of colorectal cancer (CRC), highlighting the need to identify vulnerabilities in resistant tumor cells. REV7 (MAD2L2) is a multifunctional genome-maintenance protein with established roles in REV3-dependent translesion synthesis (TLS) as well as mitotic progression, spindle organization, and chromosome segregation. Although perturbing REV7-dependent DNA damage tolerance has emerged as a strategy to overcome chemoresistance, whether pharmacologic targeting of REV7 exposes vulnerabilities to mitotic disruption remains poorly understood. Here, we investigated the REV7-targeting small-molecule inhibitor C10, which disrupts the REV7–REV3 interaction, and found that C10 sensitized HCT116 and HT-29 CRC cells to the microtubule-disrupting agent colchicine, resulting in markedly reduced clonogenic survival. In contrast, C10 did not substantially enhance sensitivity to cisplatin. Genetic loss of REV7 similarly increased sensitivity to colchicine, independently supporting a REV7-dependent vulnerability to microtubule disruption. Combined C10 and colchicine treatment was associated with pronounced alterations in mitotic and post-mitotic nuclear morphology, including condensed/mitotic-like and multinucleated configurations, and increased micronucleus formation. In HT-29 cells, combined treatment also increased the frequency of cells containing multiple γ-tubulin-positive foci, consistent with disrupted centrosome-associated organization. Together, these findings support a model in which REV7-dependent functions contribute to cellular tolerance and recovery from microtubule-mediated mitotic stress, whereas REV7–REV3 disruption renders CRC cells vulnerable to mitotic perturbation. This study broadens the potential consequences of targeting REV7 beyond its canonical role in TLS and suggests that combining REV7-directed inhibition with mitotic perturbation warrants further investigation as a therapeutic strategy in CRC and potentially other cancers.

## Introduction

REV7 (MAD2L2) is a multifunctional genome-maintenance protein best characterized as the non-catalytic regulatory subunit of DNA polymerase ζ (Pol ζ) (1). Within the four-subunit Pol ζ complex, REV7 interacts with the catalytic REV3 subunit and, together with POLD2 and POLD3, supports translesion synthesis (TLS), allowing replication to proceed across DNA lesions that would otherwise impede replicative polymerases (2). Although TLS promotes cell survival following genotoxic stress, its mutagenic nature can contribute to genome instability and the acquisition of resistance to DNA-damaging therapies (3). Consistent with these functions, elevated REV7 expression has been associated with tumor progression and therapeutic resistance across multiple malignancies (4-8), including colorectal cancer (CRC), and genetic depletion of REV7 can enhance sensitivity to DNA-damaging agents. Importantly, REV7 has been directly implicated in CRC chemoresistance: REV7 expression is elevated in drug-resistant CRC cells, while genetic disruption of REV7 restores sensitivity to 5-fluorouracil (5-FU) and oxaliplatin (9). These observations establish REV7-mediated damage tolerance as a potential therapeutic vulnerability in treatment-resistant CRC.

The therapeutic potential of targeting REV7 is further supported by the development of small-molecule inhibitors that disrupt its interactions within the TLS machinery. Actis *et al*. provided early evidence that the REV7–REV3 interface is pharmacologically tractable by identifying a small molecule that disrupts this interaction and enhances cellular sensitivity to cisplatin (10). Subsequent approaches targeting REV7-associated TLS interactions, including the REV1– REV7 interface, similarly impaired DNA damage tolerance and potentiated responses to genotoxic therapy (11). Together, these studies demonstrate that distinct protein–protein interactions involving REV7 can be pharmacologically disrupted and establish REV7 as a druggable component of the TLS machinery. Building on this foundation, we used docking-based virtual screening to identify Compound 10, a small-molecule scaffold targeting the REV7– REV3 interaction (12), and enhanced the sensitivity of mouse embryonic fibroblast (MEF) and human HT1080 cells to ionizing radiation, suggesting that the consequences of pharmacologic REV7 inhibition may extend beyond conventional TLS-associated chemotherapy responses (12).

This possibility is particularly intriguing because REV7 functions well beyond Pol ζ-mediated lesion bypass. REV7 is a core component of the Shieldin complex, which acts downstream of 53BP1–RIF1 to limit DNA-end resection and influence double-strand break repair (DSBR) pathway choice between non-homologous end joining and homologous recombination (1, 13). REV7 also directly supports cell-cycle progression and mitotic fidelity (1). It can regulate APC/C activity through CDH1 (14), associates with proteins involved in spindle organization and chromosome alignment, and loss of REV7 produces spindle abnormalities, chromosome misalignment, lagging chromosomes, anaphase bridges, and altered mitotic progression (15). These functions position REV7 at an intersection between DNA damage tolerance, DNA repair, replication completion, and chromosome segregation. Thus, while pharmacologic REV7 inhibition has been explored primarily to enhance responses to DNA-damaging therapy, whether REV7 inhibition creates a distinct vulnerability to mitotic spindle perturbation remains largely unexplored.

In this study, we tested this hypothesis using C10, a REV7-targeting small-molecule inhibitor, in HCT116 and HT-29 CRC models. We first compared the effects of REV7 inhibition in the context of DNA-damaging therapy with its effects during colchicine-induced spindle disruption (16) and then examined the cellular consequences of combined REV7 and microtubule perturbation. C10 preferentially enhanced the suppression of clonogenic survival produced by colchicine, while combined treatment produced pronounced abnormalities in nuclear and mitotic morphology that persisted following drug removal and recovery. These phenotypes included increased multinucleation and multilobulation, abnormal nuclear architecture, and alterations in centrosome-associated γ-tubulin organization, consistent with defective recovery from mitotic disruption. Together, our findings identify an unexpected vulnerability of CRC cells to combined REV7 inhibition and spindle perturbation and expand the potential therapeutic consequences of targeting REV7 beyond its established role in DNA damage tolerance.

## Methods

### Mammalian Cell Culture

HCT116 (male-derived colorectal carcinoma) and HT-29 (female-derived colorectal adenocarcinoma) cells were obtained from ATCC and maintained in McCoy’s 5A medium (ATCC) supplemented with 10% (*v/v*) fetal bovine serum (FBS; Gibco) and 1% penicillin– streptomycin (Gibco). Mouse embryonic fibroblasts (MEFs) were maintained in Dulbecco’s Modified Eagle Medium (DMEM; ATCC) supplemented with 10% (*v/v*) FBS (Gibco) and 1% penicillin–streptomycin (Gibco). Cells were maintained at 37 °C in a humidified atmosphere containing 5% CO_2_ and passaged using 0.25% trypsin.

### REV7-Dependence Assay

KP REV7 wild-type and REV7-knockout cells were seeded at 1,000 cells per well in 6-well plates in triplicate and allowed to adhere for 24 h. Cells were subsequently treated with vehicle or C10 at 2, 5, or 8 μM. Following 24 h of treatment, drug-containing medium was replaced with fresh medium, and cells were cultured for an additional 5–7 days. Colonies were fixed with 70% ethanol and stained with 1 mL of 0.5% Coomassie Blue. Colonies containing >50 cells were counted. Relative clonogenic survival was calculated by normalizing the mean colony number for each treatment condition to the corresponding vehicle-treated control for each genotype.

### Clonogenic Survival Assay

Cells were seeded at 1,000 cells per well in 6-well plates in triplicate and allowed to adhere for 24 h. Cells were subsequently treated with 5 μM C10, 25 nM colchicine (Sigma-Aldrich, C9754), cisplatin [1 μM for HCT116 cells or 5 μM for MEFs; Sigma-Aldrich, 232120], or the indicated combinations. Following 24 h of treatment, drug-containing medium was replaced with fresh medium, and cells were cultured for an additional 5–7 days. Colonies were fixed with 70% ethanol and stained with 1 mL of 0.5% Coomassie Blue. Colonies containing >50 cells were counted. Relative clonogenic survival was calculated by normalizing the mean colony number for each treatment condition to the corresponding untreated/vehicle control.

### Immunofluorescence Microscopy

HCT116 and HT-29 cells were seeded at 8,000 cells per well in 8-well chamber slides and allowed to adhere for 24 h. Cells were treated with 5 μM C10 and/or 25 nM colchicine (Sigma-Aldrich, C9754) for 24 h. Drug-containing medium was then replaced with fresh medium, and cells were allowed to recover for an additional 24 h before fixation with 4% paraformaldehyde. Cells were quenched with ammonium chloride and permeabilized with 0.1% Triton X-100 in 1× PBS. Samples were stained with antibodies against β-tubulin (Cell Signaling Technology, 86298) and γ-tubulin (Cell Signaling Technology, 5886), each at 1:100, and mounted with ProLong− Diamond Antifade Mountant and DAPI (Thermo Fisher Scientific). Samples were imaged at the University of Vermont Cancer Center Microscope Imaging Center on a Nikon A1R at 60× oil-immersion magnification.

## RESULTS

### REV7 inhibition does not broadly enhance cisplatin sensitivity

To investigate the cellular consequences of pharmacologic REV7 inhibition, we utilized Compound 10 (C10), a recently developed tetra-substituted thiophene-based small-molecule inhibitor designed to target the REV3-binding interface of REV7 and disrupt the REV7–REV3 protein–protein interaction within DNA polymerase ζ (Pol ζ) (12); **Fig. 1A**. C10 was developed through structure-guided targeting of a REV7 interaction surface centered on V150, W171, I172, D175, and D178 and inhibits the REV7–REV3-RBM1 interaction with a biochemical IC_50_ of 3.9 ± 1.0 μM (12). Although C10 was developed to disrupt the canonical REV7–REV3 interaction, whether pharmacologic perturbation of REV7 also exposes vulnerabilities associated with its non-TLS functions, including regulation of mitotic progression, remains poorly understood. We first examined the relationship between REV7 status and C10 sensitivity using clonogenic survival assays in isogenic REV7 wild-type (WT) and REV7 knockout (KO) Kras-mutant and p53-deficient (KP) cells. We exposed cells to increasing concentrations of C10 (2, 5, and 8 μM) and assessed their long-term proliferative capacity by colony formation. REV7-deficient cells retained greater clonogenic survival than REV7-WT cells at higher C10 concentrations (**Fig. 1B**), consistent with REV7 contributing to C10’s cellular activity.

**Figure 1.**
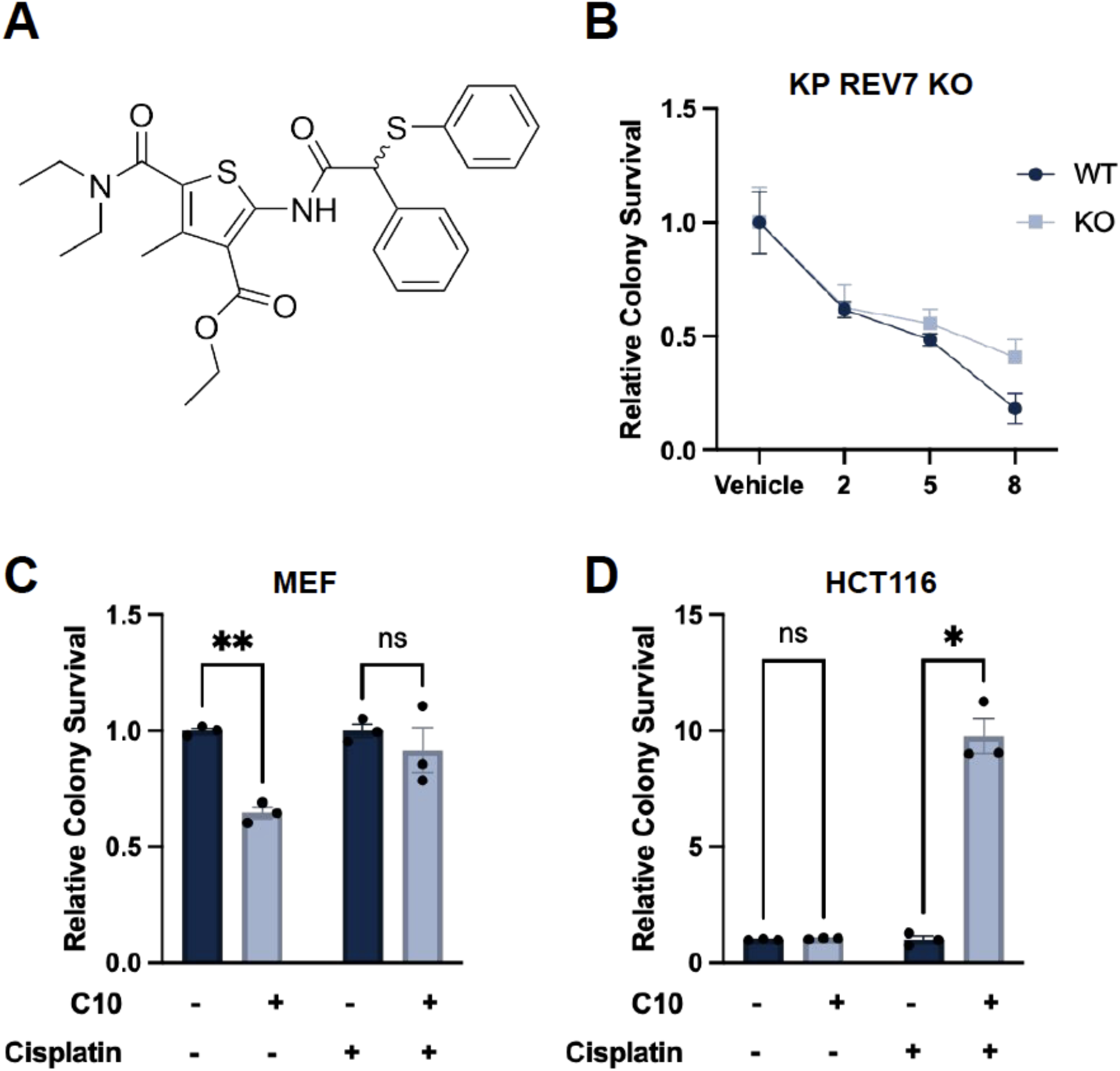
Characterization and functional specificity of the REV7-targeting compound C10. **(A)** Chemical structures of the REV7-targeting compound. **(B)** Clonogenic survival of REV7 wild-type (WT) and REV7 knockout (KO), Kras-mutant and p53-deficient (KP) cells following treatment with increasing concentrations of C10 (2, 5, and 8 µM) or vehicle. Relative colony survival was normalized to the corresponding vehicle-treated condition. **(C)** Relative colony survival of Mouse Embryonic Fibroblasts (MEF) cells following treatment with 5 µM C10 alone, 500 nM cisplatin alone, and the combination of C10 and cisplatin. **(D)** Relative colony survival of Human Colon Tumor 16 (HCT116) cells following treatment with 5 µM C10 alone, 1 µM cisplatin alone, or the combination of C10 and cisplatin. Colony survival is presented relative to the corresponding control treatment. Data are presented as mean ± S.E.M. from three independent experiments. Statistical significance was determined using multiple unpaired t-tests with Holm-Sidak correction; \****P*** < 0.05, \*\****P*** < 0.01; ns, not significant.

Because REV7-dependent translesion synthesis contributes to cellular resistance to DNA-damaging chemotherapy (2), we tested whether C10 could increase sensitivity to cisplatin. We assessed clonogenic survival in mouse embryonic fibroblasts (MEFs) and Human Colon Tumor 116 (HCT116) colorectal cancer cells after treatments with C10, cisplatin, or both. 5 µM C10 did not improve cisplatin’s (500 nM) ability to reduce clonogenic survival in MEFs (**Fig. 1C**).

Surprisingly, C10 at a 5 mM also did not make HCT116 cells more sensitive to cisplatin at 1 µM; instead, cells treated with the combination of these drugs showed higher clonogenic survival than those with cisplatin alone (**Fig. 1D**). Consistent with these findings, three structurally related substituted-thiophene REV7 inhibitors, compounds 1–3, also failed to enhance cisplatin sensitivity in HCT116 cells (**Fig. S1A–C**). These compounds share the 5-carbamoyl-substituted thiophene core, an important structural determinant of REV7 inhibitory activity, and disrupt the REV7–REV3-RBM1 interaction with biochemical IC_50_ values in the low-micromolar range (**Fig. S1D**) (12). Together, these findings suggest that pharmacologic disruption of the REV7–REV3 interaction through this compound series does not substantially sensitize HCT116 cells to cisplatin under the conditions tested.

These findings differ from previous studies in which genetic or pharmacologic perturbation of REV7 enhanced cellular sensitivity to cisplatin. In gallbladder adenocarcinoma models, REV7 knockout or knockdown increased cisplatin sensitivity, whereas re-expression of REV7 in REV7-deficient cells restored chemoresistance (5). Pharmacologic studies have similarly implicated specific REV7-containing protein interfaces in cisplatin tolerance. Actis et al. identified a small-molecule inhibitor of the REV7–REV3L interaction that bound REV7, impaired interstrand crosslink-associated DNA damage tolerance, and reduced clonogenic survival following cisplatin treatment in HeLa cells (10). More recently, pharmacologic disruption of the MAD2L2/REV7–REV1 interaction by C#3 enhanced cisplatin sensitivity across multiple cancer models, including HCT116 cells, and was associated with impaired TLS-mediated lesion bypass, reduced MAD2L2–REV1 complex formation, and persistent DNA damage signaling (11). Importantly, these approaches are not mechanistically equivalent. Genetic loss of REV7 eliminates its multiple cellular functions, whereas small molecules targeting distinct REV7 interaction surfaces may perturb different subsets of REV7-dependent activities. Moreover, the consequences of pharmacologic REV7 inhibition may depend on the targeted molecular interface, compound properties, and cellular context. Thus, our findings indicate that C10 disruption of the REV7–REV3 interface does not produce generalized sensitization to cisplatin under the conditions examined, prompting us to investigate whether C10 instead exposes vulnerabilities associated with REV7-dependent processes beyond DNA damage tolerance.

### REV7 inhibition selectively sensitizes colorectal cancer cells to mitotic spindle disruption

Beyond its canonical function in translesion synthesis, REV7/MAD2L2, has established functions in mitotic regulation (17-19). REV7 depletion disrupts mitotic spindle organization and chromosome alignment, producing abnormal spindle structures and chromosome segregation defects (15). More recently, REV3 and REV7 were shown to participate in a replication-associated checkpoint that delays mitotic entry until DNA replication is completed (20). Together, these observations raised the possibility that pharmacologic disruption of REV7 function might create a vulnerability to mitotic machinery perturbation.

We therefore examined the response to C10 in combination with colchicine, a microtubule-disrupting agent that interferes with spindle assembly and induces mitotic arrest (21, 22). We tested HCT116 and HT-29, two genetically distinct colorectal cancer (CRC) models, where HCT116 is TP53 wild-type (23), MSI-high/MMR-deficient (MLH1 and MSH3-deficient) (24), KRAS^G13D-mutant (25), and relatively chromosomally stable (26), whereas HT-29 is TP53-mutant (27), MSS/MMR-proficient, BRAF^V600E-mutant (28), and exhibits prominent chromosomal instability and aneuploidy (23, 29). These contrasting molecular backgrounds make the two cell lines complementary models for evaluating whether treatment responses are conserved across distinct CRC genomic contexts. Interestingly, in both HCT116 and HT-29 CRCs, combined C10 and colchicine treatment produced a pronounced reduction in clonogenic survival relative to either treatment alone (**Fig. 2A, B**). We next tested whether this phenotype was also genetically linked to REV7 loss. REV7-deficient KP cells showed greater sensitivity to colchicine than REV7-WT cells (**Fig. 2C**), independently supporting a role for REV7 in cellular tolerance to mitotic spindle perturbation.

**Figure 2.**
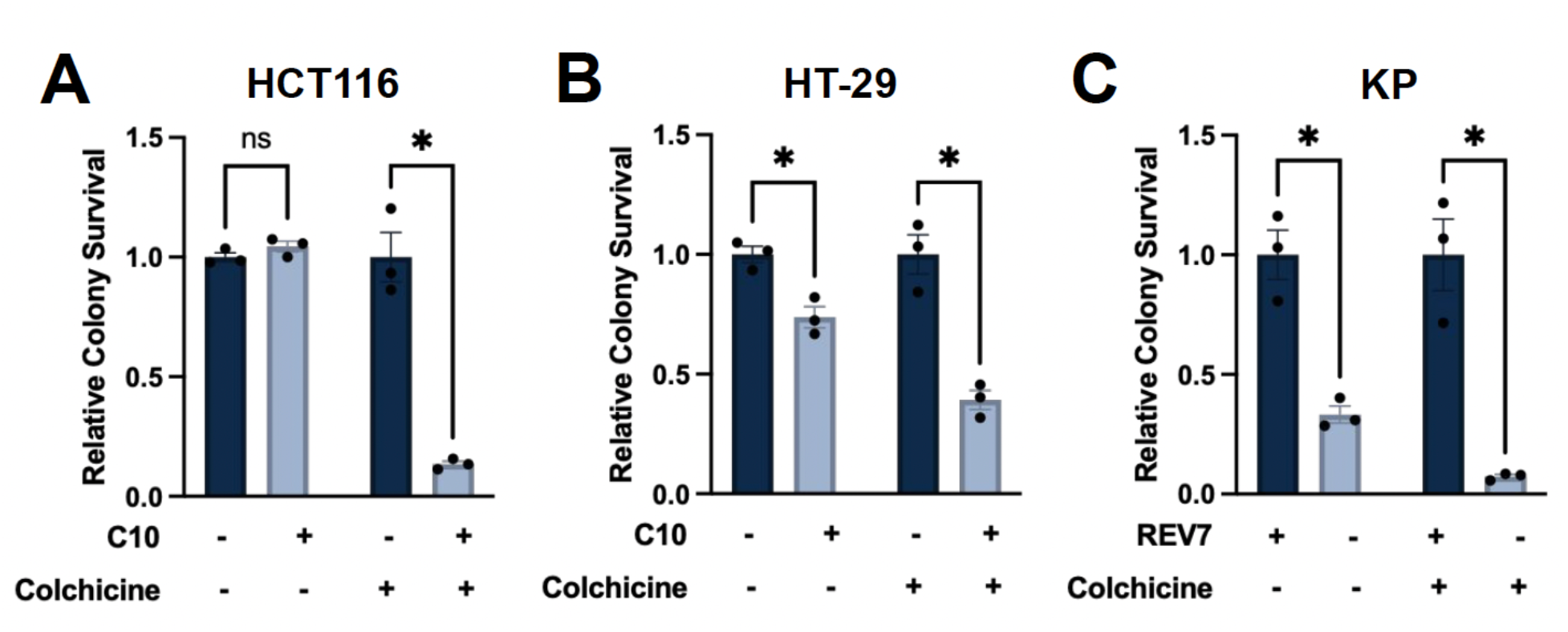
Figure 2. REV7 inhibition reduces clonogenic survival following colchicine treatment. **(A–B)** Relative colony survival of HCT116 (A) and HT-29 (B) cells following treatment with vehicle, 5 µM C10, Y 25 nM colchicine, or C10 in combination with colchicine. **(C)** Clonogenic survival of KP REV7 wild-type (REV7+) and REV7 knockout (KO) cells in the absence or presence of colchicine at 25 nM. Colony survival is presented relative to the corresponding control treatment. Data represent mean ± S.E.M. from three independent experiments. Statistical tests were performed using multiple unpaired t-tests with Holm-Sidak correction in GraphPad Prism version 11.1; \****P*** < 0.05; ns, not significant.

These results reveal an important distinction between cellular responses to DNA-damaging and mitotic stress. Although C10 did not enhance cisplatin cytotoxicity under our experimental conditions, C10 pharmacologically disrupted the REV7–REV3 interaction and markedly reduced clonogenic survival following colchicine treatment. Importantly, the increased colchicine sensitivity of REV7-deficient cells independently supports a REV7-dependent basis for this phenotype, rather than an exclusively nonspecific interaction between C10 and colchicine.

Together, these findings are consistent with REV7’s established role in spindle organization and mitotic progression and suggest that REV7-dependent functions become particularly important for cellular survival after disruption of microtubule-dependent spindle function.

### REV7 inhibition exacerbates mitotic and post-mitotic abnormalities following colchicine treatment

We next examined HCT116 and HT-29 cells by immunofluorescence to determine whether REV7 inhibition with the C10 inhibitor affects cellular mitotic stress. Cells were treated with vehicle, C10, colchicine, or the C10 + colchicine combination, recovered for 24 hours, and stained for DNA, β-tubulin, and γ-tubulin with the corresponding antibodies, as detailed in the methods (**Fig. 3A–B**). Untreated cells in both models showed predominantly intact nuclear morphology, unremarkable cellular structure, and organized microtubule architecture. DAPI staining revealed treatment-associated alterations in nuclear morphology in both HCT116 and HT-29 cells (**Fig. 3A–B; Fig. S2**). Whereas untreated cells predominantly exhibited relatively uniform nuclear morphology, C10- and colchicine-treated cells displayed irregular nuclear contours and condensed/mitotic-like chromatin. Combined C10 and colchicine treatment was associated with striking alterations in nuclear architecture, including multilobulated/multinucleated configurations and highly condensed chromatin. Qualitative differences were apparent between the two CRC models, with HCT116 cells displaying a mixture of irregular or lobulated nuclear structures and strongly condensed chromatin, whereas HT-29 cells exhibited prominent compact, highly condensed nuclear configurations.

**Figure 3.**
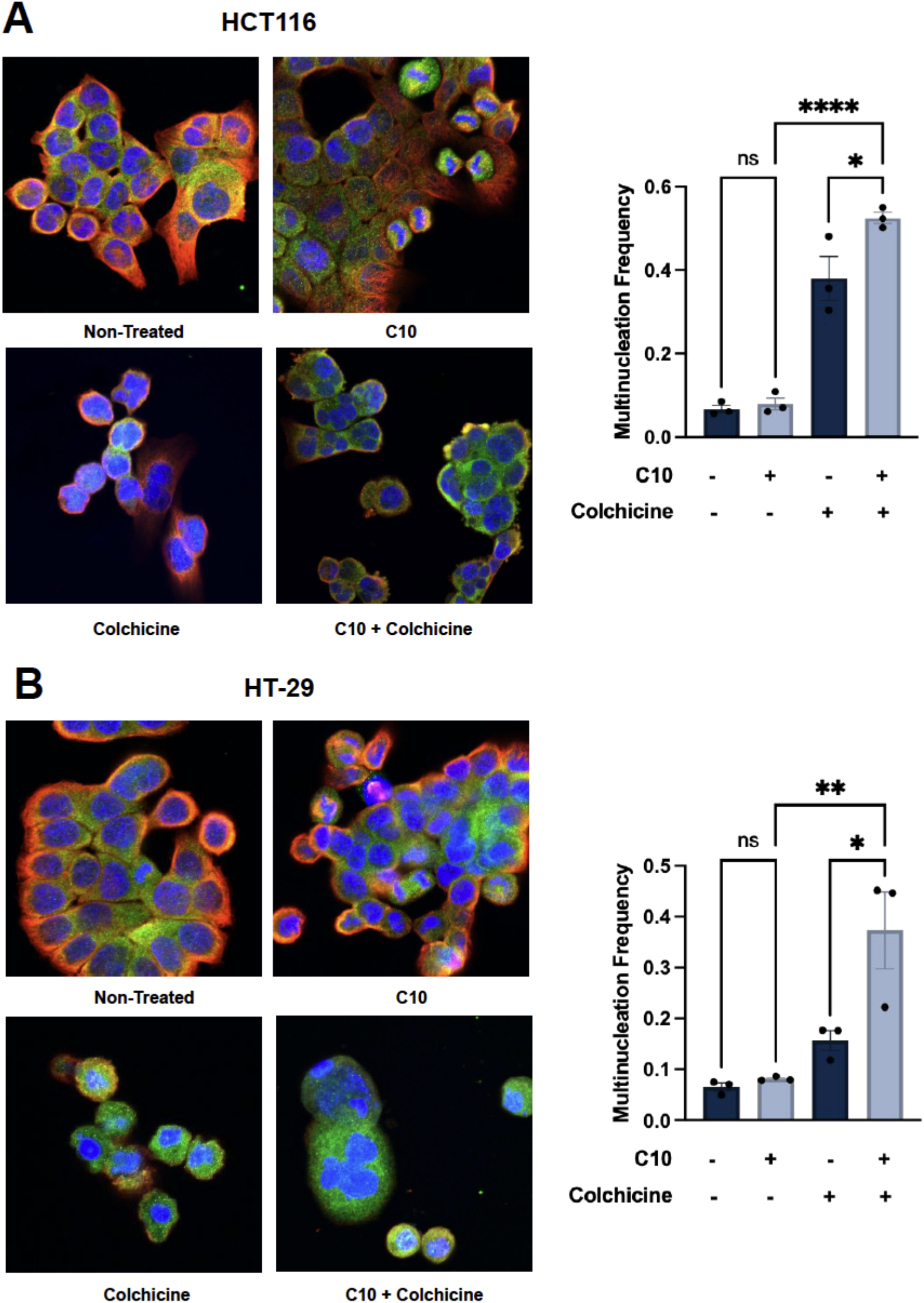
Nuclear and mitotic morphology following REV7 inhibition and colchicine treatment in CRC cells. Representative immunofluorescence images of HCT116 (top) and HT-29 (bottom) cells treated with vehicle (untreated), 5 µM C10, 25 nM colchicine, or C10 in combination with colchicine. Cells were stained for γ-tubulin (green), β-tubulin (red), and DNA (DAPI, blue). Images show representative multinucleated/multilobed cells after combined C10 and colchicine treatment. Bar graphs on the right show quantification of multinucleated/multilobed cells. Statistical significance was determined using one-way ANOVA with Tukey’s. \****P*** < 0.05, \*\****P*** < 0.01, \*\*\*\****P*** < 0.0001, Scale bar, 60 μm.

Colchicine induces mitotic stress by binding tubulin and disrupting microtubule assembly and dynamics, thereby impairing mitotic spindle formation and chromosome segregation (21). Prolonged spindle disruption can result either in mitotic cell death or in mitotic slippage, in which cells exit mitosis without completing normal chromosome segregation and cytokinesis (30). The pronounced multinucleated and multilobulated phenotypes observed after combined C10 and colchicine treatment are therefore consistent with defective mitotic progression and raise the possibility that REV7 inhibition promotes abnormal exit from colchicine-induced mitotic arrest.

This model is highly relevant because REV7 is required for functional spindle assembly and faithful chromosome segregation, and REV7 depletion produces abnormal spindles and chromosome misalignment (15). These results suggest that REV7 perturbation exacerbates the consequences of microtubule-dependent mitotic stress, potentially through impaired mitotic recovery and/or increased mitotic slippage, leading to accumulation of abnormal multinucleated cells.

DAPI staining revealed treatment-associated alterations in nuclear morphology in both HCT116 and HT-29 cells (**Fig. S2**). Whereas untreated cells predominantly exhibited relatively uniform, rounded nuclei, C10- and colchicine-treated cells displayed irregular nuclear contours and condensed chromatin. Combined C10 and colchicine treatment was associated with striking alterations in nuclear architecture, including multilobulated/multinucleated configurations and highly condensed chromatin. Qualitative differences were also apparent between the two cell lines, with HCT116 cells displaying a mixture of irregular or lobulated nuclear structures and strongly condensed chromatin, whereas HT-29 cells exhibited prominent compact, highly condensed nuclear configurations. These phenotypes are consistent with defective mitotic or post-mitotic outcomes and suggest that REV7 inhibition compromises cellular recovery from colchicine-induced spindle stress. Prolonged spindle disruption can result in aberrant mitotic exit, including mitotic slippage, in which cells escape mitotic arrest without completing normal chromosome segregation and cytokinesis (30). Thus, the multilobulated/multinucleated configurations observed following combined treatment are compatible with aberrant mitotic exit, potentially involving mitotic slippage and/or cytokinesis failure. This is particularly intriguing given REV7’s established role in spindle organization and chromosome segregation (15). However, because these experiments capture fixed-cell morphology, these images alone cannot resolve the precise mitotic events that give rise to these phenotypes.

Because aberrant mitotic exit and chromosome segregation defects can result in micronuclei, we next asked whether the pronounced nuclear abnormalities observed after combined REV7 inhibition and microtubule disruption were accompanied by increased micronucleus formation. Micronuclei were readily detected after colchicine treatment and were particularly prominent after combined C10 and colchicine treatment in both HCT116 and HT-29 cells (**Fig. 4**). Quantification showed a significant increase in micronucleus frequency with the combination compared with either C10 or colchicine alone in both the cell lines, with the effect slightly more pronounced in HCT116 cells. These findings provide additional evidence that REV7 inhibition exacerbates the consequences of microtubule disruption and are consistent with increased chromosome segregation errors and/or defective resolution of mitotic stress.

**Figure 4.**
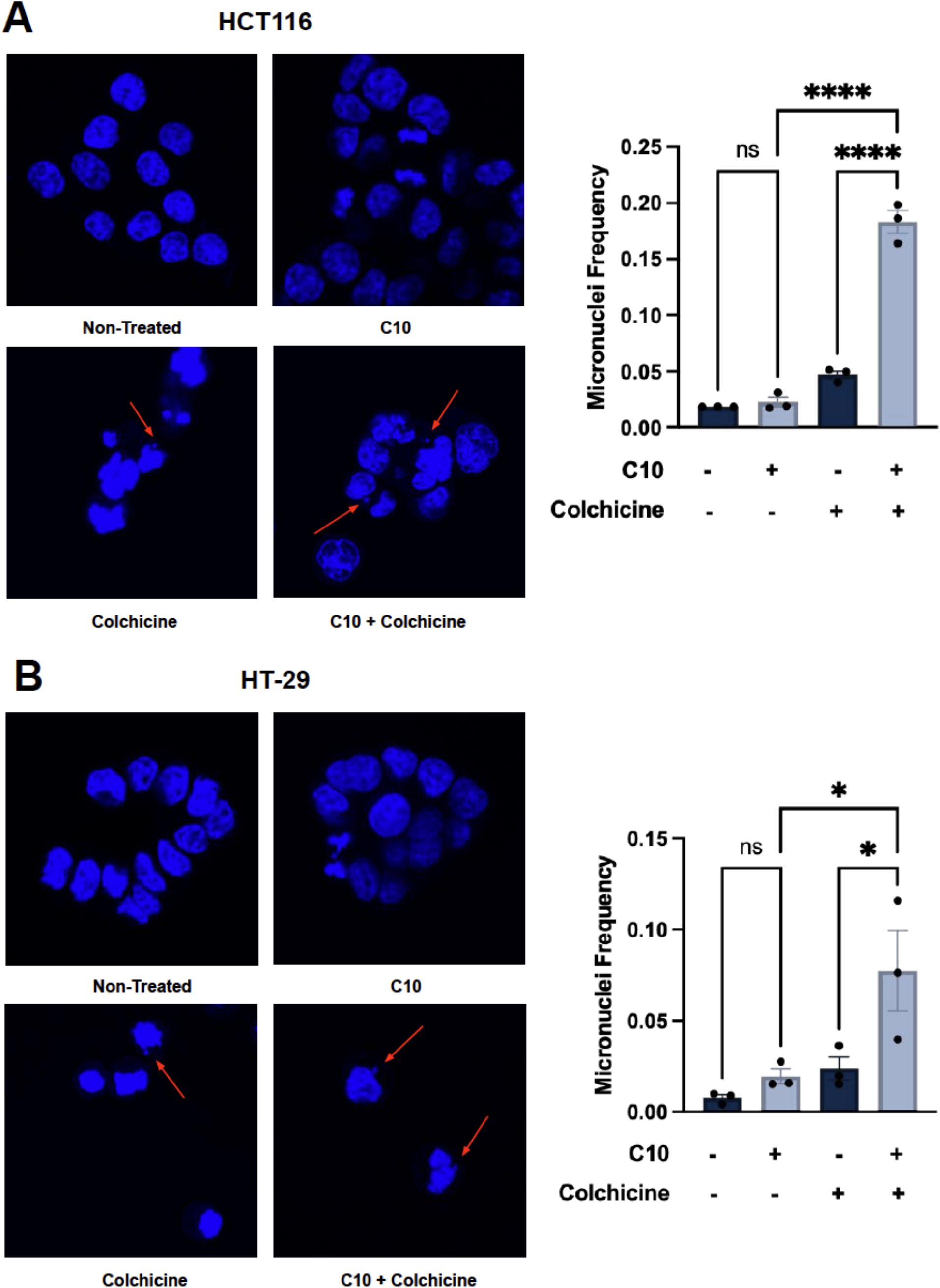
Micronucleus formation following REV7 inhibition and colchicine treatment in CRC cells. Representative DAPI-stained images of **HCT116 (A)** and **HT-29 (B)** cells following treatment with vehicle (non-treated), C10, colchicine, or C10 plus colchicine. DAPI staining (blue) visualized DNA, and red arrowheads indicate representative micronuclei. Right, quantification of micronuclei following the indicated treatments, expressed as the percentage of cells containing micronuclei. Statistical significance was determined using one-way ANOVA with Tukey’s post hoc test. Data are from 3 independent experiments with approximately 100 cells analyzed per condition per experiment. Error bars represent mean ± S.E.M. Scale bar, 60 μm. \****P*** < 0.05, \*\*\*\****P*** < 0.0001.

Although micronucleus formation was particularly pronounced in HCT116 cells, the comparatively modest micronucleus response in HT-29 cells suggested that mitotic disruption might manifest differently across CRC backgrounds. We therefore examined centrosome organization in HT-29 cells using γ-tubulin as a marker of centrosome-associated microtubule-organizing centers. Whereas untreated, C10-treated, and colchicine-treated cells predominantly displayed one to two discrete γ-tubulin foci, combined C10 and colchicine treatment produced cells with multiple γ-tubulin-positive foci (**Fig. 5**). Quantification showed a significant increase in the frequency of cells with >2 γ-tubulin foci after combined treatment compared with either treatment alone. Together with the micronucleus phenotype, these findings indicate that combined REV7 inhibition and microtubule disruption perturb multiple features of mitotic fidelity, with the predominant manifestation differing between CRC cell backgrounds. The emergence of multiple γ-tubulin-positive foci in HT-29 cells is consistent with abnormal centrosome organization and provides an additional potential mechanism through which combined treatment could promote defective spindle organization and chromosome segregation. A limitation of this analysis is that γ-tubulin staining alone cannot distinguish true centrosome amplification from fragmentation or redistribution of pericentriolar material. Thus, the increased number of γ-tubulin-positive foci should be interpreted as abnormal centrosome-associated organization, and confirming bona fide centrosome amplification requires additional centriole/centrosome markers.

**Figure 5.**
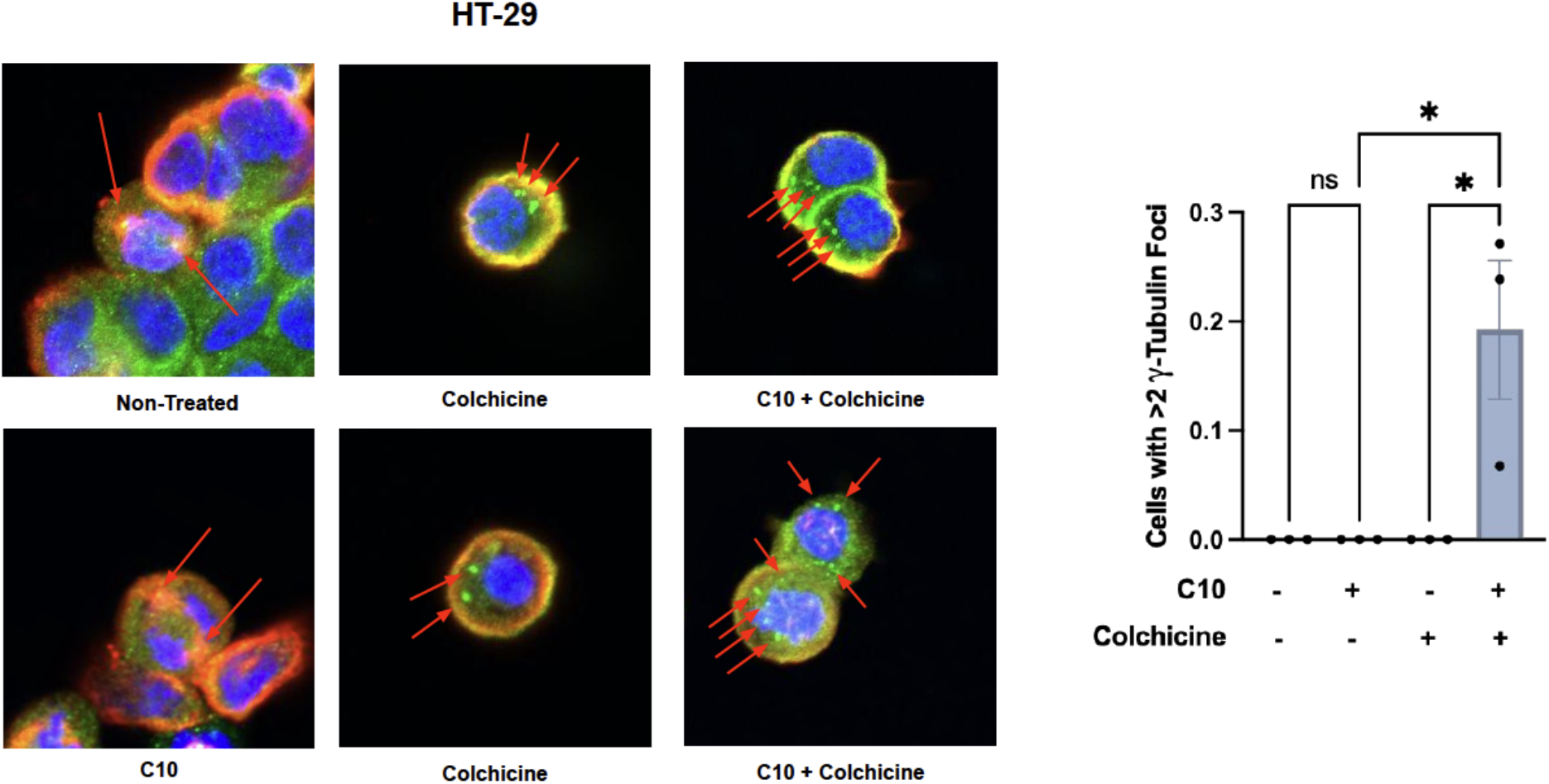
Combined C10 and colchicine treatment alters centrosome-associated γ-tubulin organization in HT-29 cells. Representative immunofluorescence images of HT-29 cells following treatment with vehicle (non-treated), C10, colchicine, or C10 in combination with colchicine. Cells were stained for γ-tubulin (green), β-tubulin (red), and DNA with DAPI (blue). Arrows indicate discrete γ-tubulin-positive foci. Quantification shows the frequency of cells containing >2 γ-tubulin-positive foci under each treatment condition. Data represent 3 independent experiments and are presented as mean ± S.E.M. Statistical significance was determined using one-way ANOVA with Tukey’s; \****P*** < 0.05; ns, not significant. Scale bar, 60 μm.

Centrosomes function as the major microtubule-organizing centers of animal cells and are critical for bipolar spindle assembly and faithful chromosome segregation (31). Numerical or structural centrosome abnormalities can perturb spindle organization and increase chromosome segregation errors, thereby contributing to chromosomal instability (32, 33). In CRC specifically, centrosome amplification and instability have been associated with aneuploidy and numerical chromosomal abnormalities (34, 35). Thus, the increased frequency of cells with >2 γ-tubulin-positive foci after combined C10 and colchicine treatment is consistent with disrupted centrosome-associated organization and further indicates impaired mitotic fidelity. This phenotype may be mechanistically related to the chromosome-segregation defects reflected by micronucleus formation, as centrosome abnormalities can promote erroneous chromosome segregation and micronucleus-generating events (36, 37). However, γ-tubulin primarily identifies pericentriolar material and microtubule-organizing centers, and γ-tubulin staining alone does not establish that individual foci represent structurally intact centrosomes containing supernumerary centrioles (38-40). Accordingly, confirmation of bona fide centrosome or centriole amplification will require additional centrosomal/centriole markers (41, 42).

Collectively, these findings support a model in which REV7-dependent functions contribute to CRC cells’ ability to tolerate and recover from microtubule-mediated mitotic stress. We therefore propose that combined disruption of the REV7–REV3 interaction and microtubule dynamics compromises mitotic recovery, resulting in abnormal mitotic and post-mitotic outcomes and ultimately reduced clonogenic survival (**Fig. 6**). In this model, colchicine-induced microtubule disruption creates mitotic stress that becomes particularly detrimental when the REV7–REV3 interaction is simultaneously disrupted by C10. Combined treatment was associated with abnormal nuclear architecture, multinucleated/multilobulated configurations, micronucleus formation, and altered centrosome-associated γ-tubulin organization, together with a pronounced reduction in clonogenic survival. Although the precise sequence of events underlying these phenotypes remains to be established, our findings identify mitotic stress as a vulnerability associated with pharmacologic disruption of REV7–REV3 in CRC cells and distinguish this response from the limited sensitization observed with cisplatin under the conditions examined. This study broadens the potential consequences of targeting REV7 beyond its canonical role in DNA damage tolerance and suggests that combining REV7-directed inhibition with mitotic perturbation warrants further investigation as a therapeutic strategy in CRC, and potentially other cancers.

**Figure 6.**
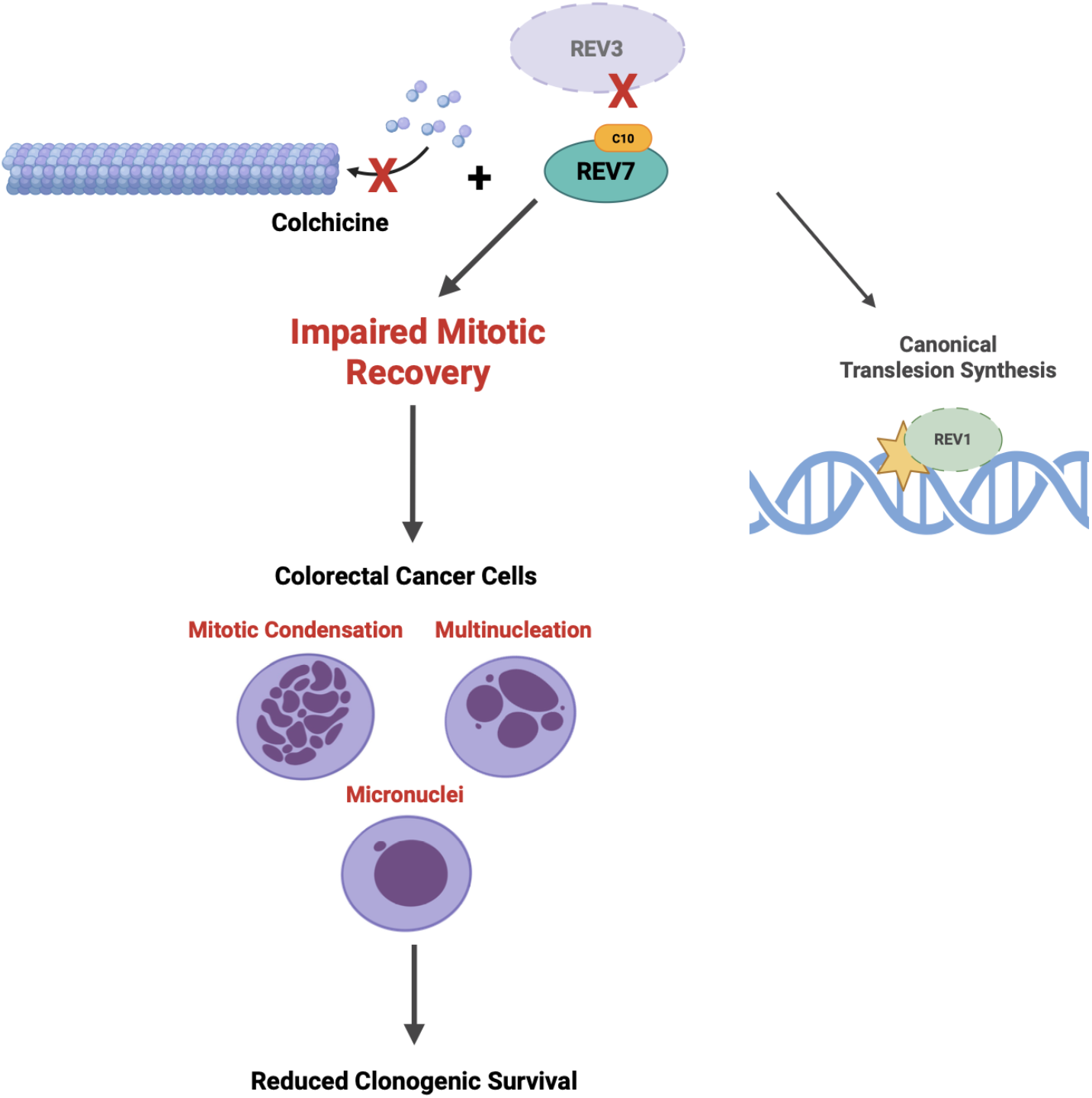
Proposed model for REV7-dependent tolerance of mitotic stress in colorectal cancer cells. REV7 participates in canonical REV3-dependent translesion synthesis but is also implicated in regulating mitotic progression and spindle function. In the proposed model, colchicine-mediated disruption of microtubule dynamics combined with C10-mediated disruption of the REV7–REV3 interaction compromises cellular recovery from mitotic stress. Combined treatment is associated with abnormal mitotic/post-mitotic nuclear morphology, including condensed/mitotic-like and multinucleated configurations, micronucleus formation, altered centrosome-associated γ-tubulin organization, and reduced clonogenic survival. The precise mechanisms connecting REV7–REV3 disruption to these mitotic phenotypes remain to be established. The model summarizes the experimental findings and proposes a mechanistic interpretation of this study.

## Author Contributions

M.G., N.K., V.G., and A.C. clonogenic survival assays; M.G., S.B., and K.B. immunofluorescence microscopy; N.H. drug synthesis and validation; M.G. and K.B. figures and data tabulation; J.R. data and statistical analysis; D.K. and M.K.H. drug synthesis and supervision; N.C. conceived the project, designed and supervised the experiments, and wrote the paper with the help of all authors.

## Acknowledgements

We thank all members of the Chatterjee Lab for contributing to this project. We also thank the University of Vermont Cancer Center (UVMCC) for the general support with research infrastructure. This study is supported by a National Cancer Institute Grant R01CA233959 to M.K.H and D.M.K., the 2026 UVMCC Pilot Project, the NIGMS R35GM150992 to N.C., and the Nicole J. Ferland and Distinguished Undergraduate Summer Research Awards from the UVM Department of Microbiology & Molecular Genetics.

