## Supplemental Material for "Targeting REV7 exposes a mitotic vulnerability in colorectal cancer cells"

**Supplementary Material**

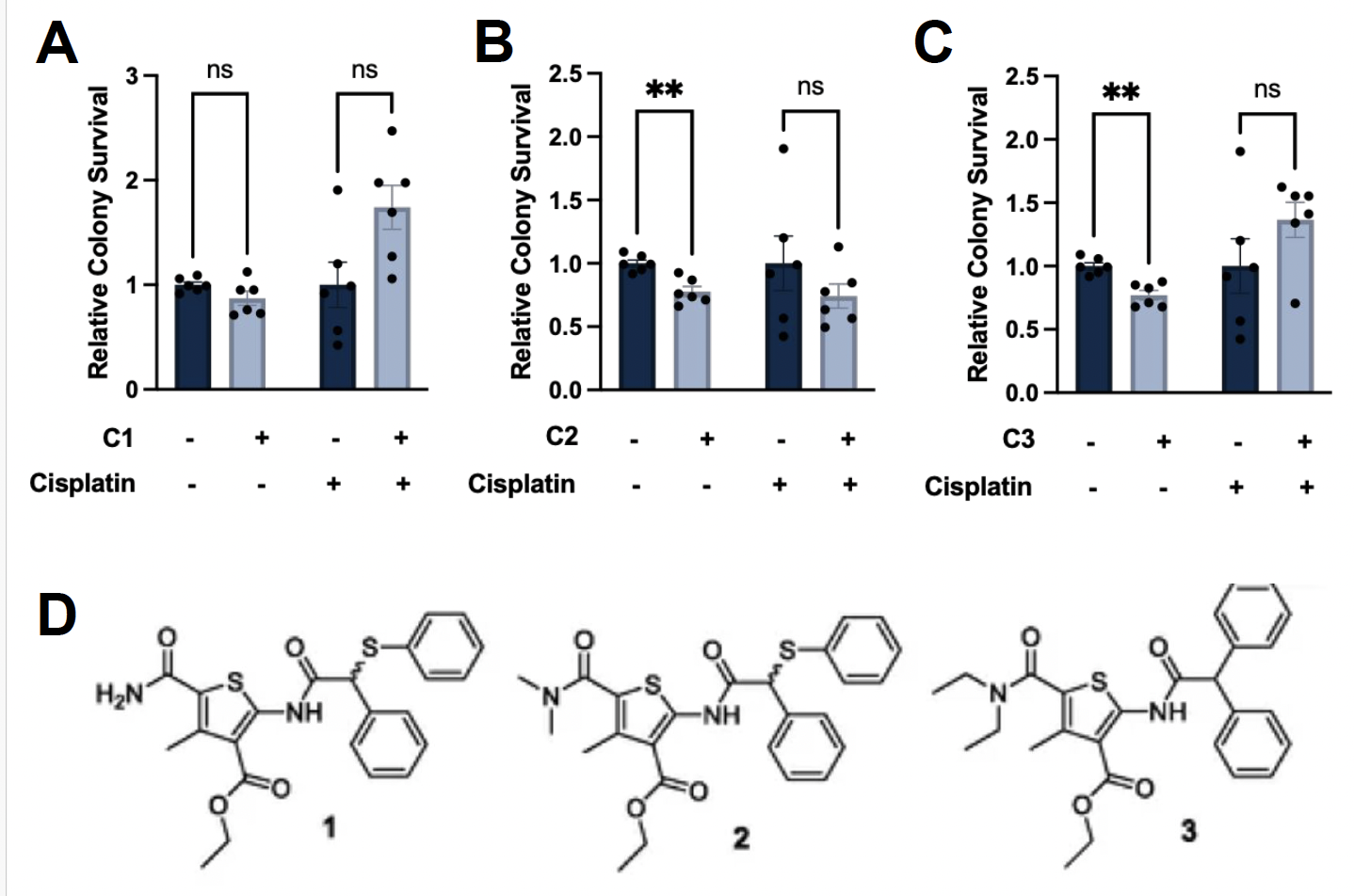

**Figure S1. Effects of REV7-targeting compounds on clonogenic survival of HCT116 cells in combination with cisplatin.** **(A–C)** Clonogenic survival of HCT116 cells after treatment with the REV7-targeting compounds C1 **(A)**, C2 **(B)**, or C3 **at 10** µM, with or without cisplatin (1 µM). Colony survival is presented relative to the corresponding untreated control. Data represent mean ± S.E.M., n=6. Statistical tests were performed using multiple unpaired t-tests with Holm-Sidak correction; ***P*** < 0.01; ns, not significant. **(D)** Chemical structures of compounds 1–3, structurally related substituted-thiophene inhibitors of the REV7–REV3-RBM1 protein–protein interaction. The compounds share a 5-carbamoyl-substituted thiophene core, an important determinant of activity in this series. Biochemical IC₅₀ values for disruption of the REV7–REV3-RBM1 interaction were 11.5 ± 4.9 μM, 10.4 ± 1.1 μM, and 4.8 ± 1.5 μM for synthesized compounds 1, 2, and 3, respectively (Harrahill et al., 2026).

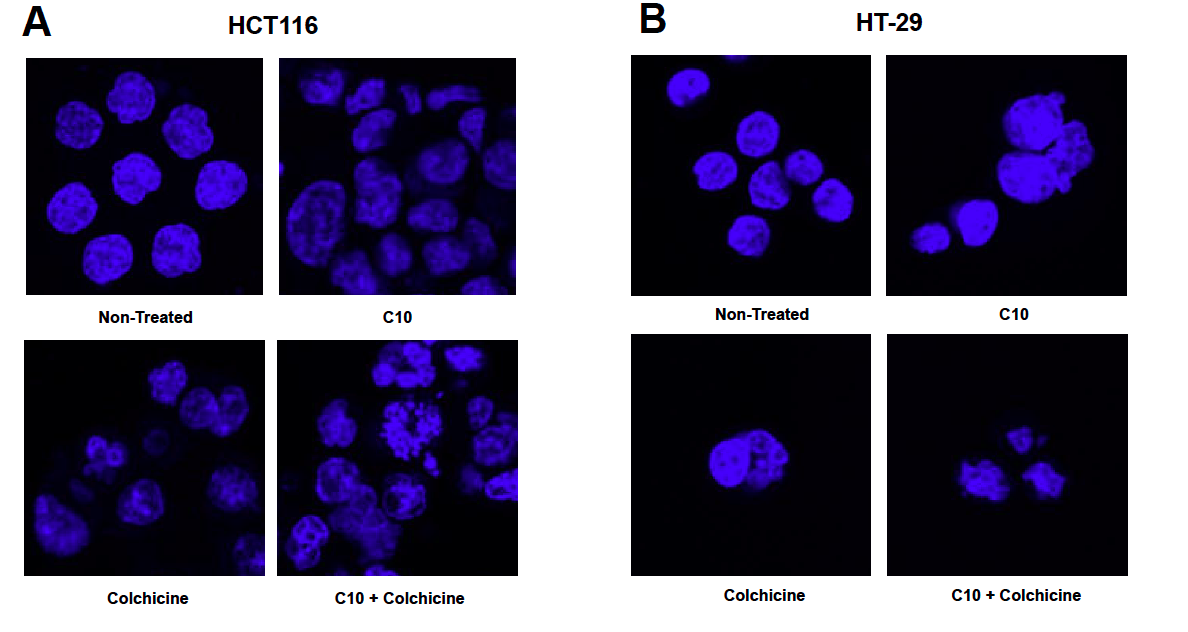
**Figure S2. Nuclear morphology following C10 and colchicine treatment in colorectal cancer cells.** Representative DAPI images of **HCT116 (A)** and **HT-29 (B)** cells following treatment with vehicle (non-treated), C10, colchicine, or C10 in combination with colchicine. DAPI staining (blue) visualized DNA. Images are representative of 3 independent experiments. Scale bar, 60 μm. Given the heterogeneous nature of the observed nuclear phenotypes and the absence of a predefined objective classification scheme, nuclear morphology was not quantitatively scored.
